# Phenological turnover matters when making trait-based predictions of plant-pollinator interactions

**DOI:** 10.1101/2025.01.10.632321

**Authors:** Aoife Cantwell-Jones, Juliet Everson, Olivia K. Bates, Abdullah M. R. Al-Hayali, George Allen, Lucas Berard, Frauke Caliebe, Suzannah Egleston, Lucia Hudson, Jacqui A. S. James, Lena Jung, Moganavalli Kattan, Renée Lejeune, Johan Svedin, Silas van Brakel, Manon van Unen, Jason M. Tylianakis, Keith Larson, Richard J. Gill

## Abstract

1. Understanding the processes determining species’ interactions is key to predicting and safeguarding ecological networks under rapid environmental change. One approach to estimating interactions is to use morphologies of taxa interacting across trophic levels to reveal suites of traits they are more likely to interact with (i.e. a trait niche).
2. Previous work studying these morphological trait niches has typically used interactions between species that are pooled in space and time. However, species assemblages, and the traits of individuals within species, can change across even small landscapes over a season, leading to morphological trait space being dynamically reshaped. Therefore, it is unclear how morphological trait turnover affects our inferences of trait niches, and our ability to answer this is in part limited by a lack of individual-level trait data.
3. Here, we directly address this by studying a montane Arctic plant-pollinator community over five growing seasons (>1,300 hours of fieldwork). Specifically, we linked every recorded plant-bumblebee interaction with the traits of the bee individual involved (n = 1,150), to investigate 1) whether plant taxa (n = 10) exhibited bee trait niches by interacting with specific regions of multidimensional trait space of visiting bumblebees, and 2) how our inference of these trait niches was affected by considering bumblebee trait turnover and plant taxon turnover.
4. When not considering turnover (interactions in space and time are pooled), plant taxa demonstrated bee trait niches. However, next we considered how bee trait space is reshaped over the elevational and seasonal gradient (especially with the emergence of different castes), and how this reshaping co-occurs with different spatiotemporal ranges of the plant taxa. From this we found the degree to which plant taxa exhibited trait niches declined significantly, and that seasonal reshaping of bee trait space was the primary driver of this trend.
5. Overall, in highly dynamic systems, like the Arctic, overlooking community turnover could mask and even overestimate the ability of morphology to explain interactions. Hence, determining how morphological traits of individual interaction partners are in phenological synchrony at localised scales will be fundamental to understanding the role morphology plays in underpinning plant-pollinator interactions.

## 1. INTRODUCTION

Understanding the mechanisms that determine species’ interactions is key to predicting ecological network responses to environmental change (Cantwell-Jones et al., 2024; Schleuning et al., 2020), and forecasting risks to ecosystem function and productivity (Schleuning et al., 2015; Tylianakis & Morris, 2017). Trait-based approaches are being increasingly used to predict interactions (e.g. Pichler et al., 2020), such as using morphology to determine the likelihood of two species that co-occur interacting (Bender et al., 2018; Garibaldi et al., 2015). For instance, gape size of frugivorous birds can covary with fruit shape (Grant & Grant, 2006), as can pollinator proboscis length with flower corolla depth (Johnson & Anderson, 2010; Stang et al., 2009). The likelihood of two species interacting at any given time or place, however, is unlikely to be determined by just a single trait. Instead, multiple morphological traits are needed to overcome acute environmental constraints (e.g. body and wing size mediating temperature-dependent activity; Kenna et al., 2021) before an interaction can occur. Therefore, while a single trait may explain interactions patterns between species across whole seasons or geographical ranges, a more multidimensional trait-based approach may be needed to dynamically predict interactions at more localised spatiotemporal scales (CaraDonna et al., 2017).

To date, most studies pool interactions across space and time, due often to low sampling resolution (Dehling et al., 2016; Eklöf et al., 2013; Peralta et al., 2023; Pichler et al., 2020, but see Liang et al., 2021; Sponsler et al., 2022). Yet we know interactions occur between individuals at small spatial scales and over short timeframes (Arroyo-Correa et al., 2023), and that morphological traits commonly turn over across landscapes and over the season within populations and species (Cantwell-Jones et al., 2023; Kalske et al., 2012), and different community compositions (Classen et al., 2017; de la Riva et al., 2018; Zhang et al., 2023). For example, individual plants may produce flowers with different phenotypes between spring and summer (Gómez et al., 2020), while environmental filtering may shape the phenotypes available at a given location (e.g. filtering out of small-bodied individuals within species at high elevation; Classen et al., 2017). More broadly, trait-environment relationships (such as, with temperature or aridity) can lead to inter- and intraspecific trait turnover over local gradients (Kemppinen & Niittynen, 2022; Lambrecht & Dawson, 2007; de la Riva et al., 2018; Rixen et al., 2022). Such turnover means that the multiple morphological dimensions that a given species can interact with can dynamically change (Klomberg et al., 2022). Yet, to our knowledge, explicit datasets enabling such trait turnover to be linked with interaction networks are almost non-existent. This has limited our ability to map individual-level data in multidimensional space to assess species’ trait preferences (but see Arroyo-Correa et al., 2021; Crestani et al., 2019; Nagano et al., 2014; Vissoto et al., 2022), confining us to assign single and averaged trait values to each species (Cantwell-Jones et al., 2024). Empirical studies are thus urgently needed to collect data where individual-level interactions are associated with multiple individual-specific morphological traits at a high spatiotemporal resolution, i.e. data that encompass natural spatial and seasonal gradients.

Here, we investigated whether Arctic plant taxa of a localised community show trait niches by interacting with a subset of pollinator (bumblebee) morphological trait space, and how phenological and elevational community turnover impacted our detection of trait niches. This was only achievable by having an individual-level plant-pollinator interaction dataset, which we collected over a five-year period at plots sampled on average twice per week, every week, for 2-3 months. By measuring multiple traits of each bumblebee individual per interaction (intertegular distance [distance between forewing attachment points], wing length, head width and estimated proboscis length), we were able to represent each individual bumblebee pollinator in multidimensional trait space (in which each axis represents a combination of traits, and a bumblebee’s position corresponds to its trait values; Hutchinson, 1957) and map trait availability in space and time. This allowed us to characterise the trait niche of a given species by calculating the volume of its interaction partners’ trait space (Dehling et al., 2016; Junker et al., 2013). We hypothesised that if bumblebee morphology is a strong predictor of plant-bumblebee interactions (e.g. Liang et al., 2021; Sponsler et al., 2022), we should expect each plant species to interact with a morphological subset of the bumblebee trait space (Fig. 1). As bumblebees show large intraspecific variation due to the presence of castes (i.e. behaviourally and morphologically distinct classes of individuals within social insect species, here queens, workers and drones), two individuals from different species can be more morphologically similar (e.g. in body size) than two individuals from the same species (Cantwell-Jones et al., 2023; Methods S1; Fig. S1-3; Table S1). We therefore focused on the traits of the individual bumblebees that plants interacted with (mixing intra- and interspecific trait variation) and ignored species identity.

**Figure 1:**
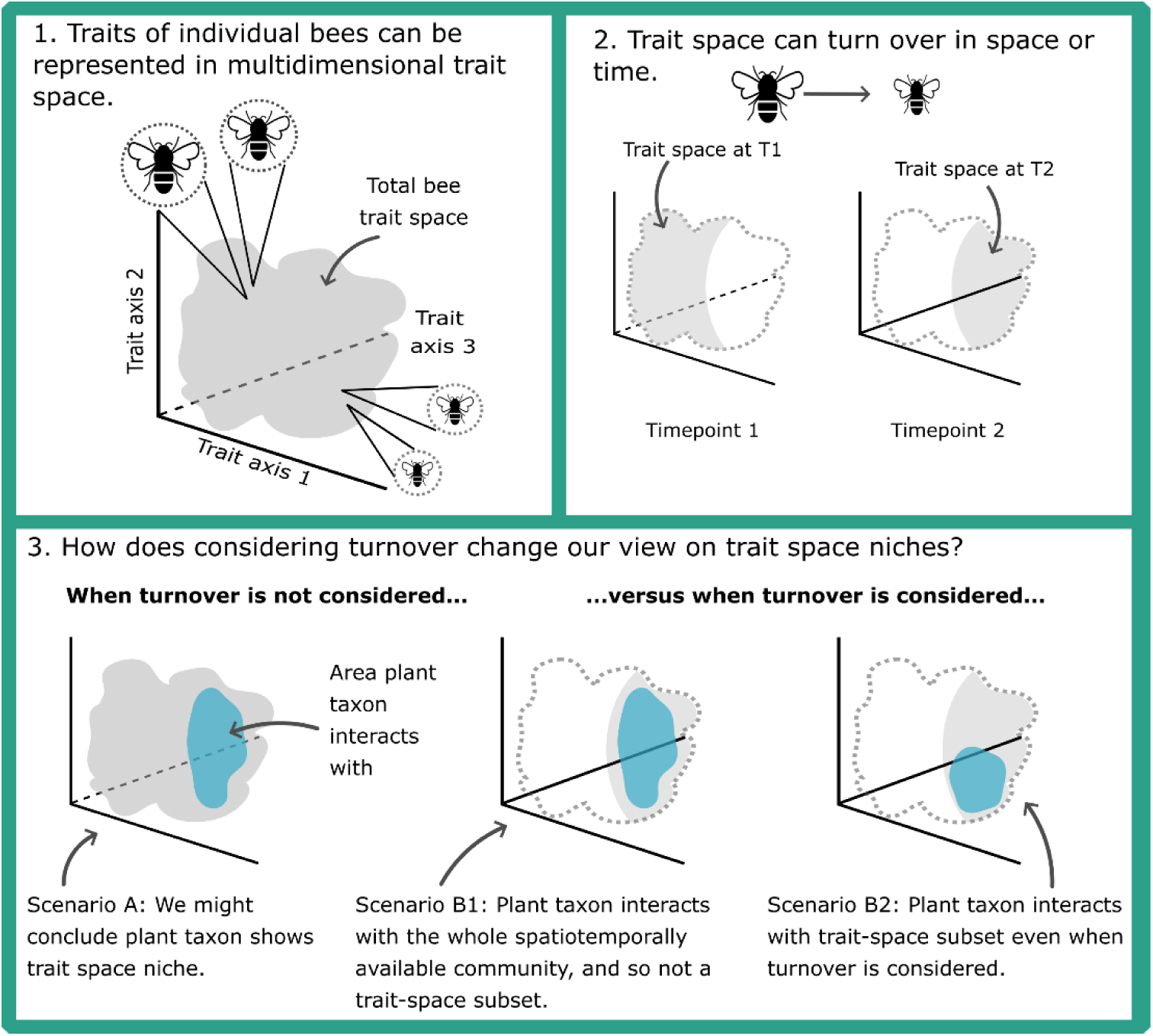
Conceptual representation of the key questions. (1) Represents multidimensional trait space, with the grey polygon representing the community of individual bees dispersed across the trait space to create a hypervolume. (2) Highlights how the trait space of a community could change over time (or space). In our study system, this could occur due to the emergence of different castes within bee species, leading to large intraspecific turnover without species turnover. (3) Illustrates the different hypotheses about how trait turnover could affect signals of “trait niches”, where plant taxa show a trait niche by interacting with a specific region (in blue) of the bumblebee-community trait space. Scenario A suggests that when turnover is not considered, we might infer that the plant taxon has a trait niche by interacting with a subset of the total, spatiotemporally-pooled bee trait space. Scenario B1 posits that the niche observed in Scenario A is potentially misleading, as when localised turnover is considered, a trait niche is not observed relative to the spatiotemporally available community (in grey). Scenario B2 posits that the plant taxon could still show a trait niche, by interacting with a subset of bee trait space even when the spatiotemporally available bumblebee community is accounted for.

Specifically, we question: 1) do plant taxa interact with specific regions of bumblebee community trait space, i.e. a morphological subset that has a smaller volume than the trait space of the whole bee community, which we refer to as a plant species’ trait niche? Then, we ask: 2) are our inferences of plant species trait niches impacted by considering turnover of the whole community, i.e. changes in actual co-occurrence of bees and plants? We tested this by comparing niches when traits were considered over the entire collection season and elevation range (ignoring community turnover and co-occurrence; Fig. 1) to the niches considering the spatiotemporally available trait space (i.e. actual co-occurrences between individual bees and plant taxa). Our system is ideal to study the aforementioned questions given: 1) the short Arctic growing season allows us to capture the community dynamics within a few months, 2) there is a relatively low turnover of bee species over time but high (intraspecific) turnover of traits (Methods S1; Figs. S1-3; Tables S1-3) as the different castes emerge, 3) the harsh climatic conditions likely impose levels of environmental filtering on traits available at specific points in space (elevational) and time (seasonal).

## 2. MATERIALS & METHODS

### 2.1 Plant-pollinator system and study site

We collected plant-bumblebee interaction data across 13 permanent 45 × 45 m plots located along a 3.4 km transect spanning 418-1164 m.a.s.l (Fries, 1925) up the eastern face of Mt. Nuolja, Abisko National Park, Sweden (68° 22’ N, 18° 47’ E). We conducted observations over five seasons (2018: 24^th^ May – 20^th^ July; 2019: 13^th^ May – 19^th^ July; 2021: 22^nd^ May – 29^th^ July; 2022: 22^nd^ May – 29^th^ July; 2023: 18^th^ May – 31^st^ July), with each permanent plot visited a mean 2.02 times per week (± standard deviation: 0.65; Fig S4, Table S4). To survey plant-bumblebee interactions within each plot, we walked a set route with sampling effort standardised to 20 min (citation removed for author anonymity). We recorded all bumblebee species observed foraging inside the plot (*n* = 3,404 foraging observations; 17 taxa) and identified the plant species they were visiting (*total* = 41 plant taxa). Exceptions to this were species belonging to the genera *Salix*, *Taraxacum*, *Myosotis* and *Cirsium*, for which we respectively grouped species to genus level, due to either hybridisation among congeners (*Salix*), or difficulty identifying to species level (*Taraxacum, Myosotis* and *Cirsium*). We focused on the bumblebee community to 1) be able to collect high-resolution, individual-level data on an entire pollinator group, and 2) as they are the first pollinators to become active in the Arctic, visiting flowers even when there is high snow cover (Bergman et al., 1996; Kevan, 1973; Lundberg & Ranta, 1980).

For bumblebee-trait data collection, we attempted to catch each observed bumblebee with a butterfly net (40 cm diameter). When successful, we transferred each caught bumblebee into an individual, lidded plastic holding-pot (150 mL) and placed them into a dark, insulated bag. At the end of each survey, and in order of capture, we transferred caught bees to a marking cage (with scale bar) and took a top-down dorsal image using a digital camera (Canon SX720 HS or Samsung Galaxy A21s), from which we measured bee morphological traits (Section 2.3). We non-lethally identified bumblebees to species level in the field based on morphological features (following Söderström, 2017; except the cryptic species *B. alpinus* and *B. polaris* that we grouped to *B. alpinus/polaris*, and *B. norvegicus* and *B. sylvestris* that we grouped to *B. norvegicus/sylvestris*; note *B. polaris* in Sweden has recently been assigned *B. pyrrhopygus*; Williams et al., 2019). We assigned identifications a confidence score between one and three (with three being most confident). We only included species in the analysis with confidence scores greater than or equal to two (n = 2,846 bees). We note that each of the studied bumblebee species was active during the majority of the seasonal period and elevational gradient we analysed (Fig. S5-6). Two exceptions to this were *B. pascuorum* and *B. alpinus/polaris*, which tended to be found at relatively lower (434.4-901.1 m.a.s.l) and higher (582.6-1159 m.a.s.l) elevations, respectively.

### 2.2 Determining spatiotemporal ranges of each plant taxon

Plant spatiotemporal data were collected by the ongoing Nuolja phenology project, in which plant developmental stages (phenophases) were recorded every five days on average from May 25^th^ until the end of the season (2017-2023) for each of the 78 segments (ca. 45 m in length) constituting the length of the Mt. Nuolja transect (MacDougall et al., 2021). We analysed the phenophases related to flowering, given this is the structural feature that determines a pollinator interaction. To estimate the flowering phenological windows for each of our bumblebee-visited plant taxa, we used the range of days (first and last dates) each taxon was observed flowering each year. Similarly, to estimate the elevational distribution, we used the elevational range (lowest and highest elevation, based on presence/absence of a taxon flowering in a segment) that each taxon was recorded in for each year. As range can be affected by outliers and to ensure conservative comparative analyses, we removed the first and last 5% of data points of each taxon’s phenological and elevational recordings each year (Fig. S7-8). These distributions were used to determine which individual bumblebees each plant taxon co-occurs with.

### 2.3 Measuring bumblebee morphology

We collected individual-level data on four morphological traits for bees: intertegular distance (ITD; distance between forewing attachments to the thorax; Cane, 1987), head width (i.e. the largest distance between distal surfaces of the eyes measured dorsally; del Castillo & Fairbairn, 2012); wing length (maximal distance between one forewing attachment point and the most distal point on that corresponding wing’s marginal cell); and estimated proboscis length (length of prementum) (Fig. 2). ITD is related to foraging performance (Klein et al., 2017), thermoregulatory ability (Bishop & Armbruster, 1999), and ability to fly at low temperatures (Kenna et al., 2021). Head width, wing length and proboscis length are related to, respectively, learning ability (Gronenberg & Couvillon, 2010), flight efficiency (Dillon et al., 2006), and floral compatibility (i.e. trait matching; Liang et al., 2021; Ranta & Lundberg, 1980)(Table S5). We measured ITD, wing length and head width from the images taken of bumblebees in the field, by landmarking images using TPS software (tpsDig v. 2.3.2 & tpsUtil v. 3.2; Rohlf, 2015) and exporting the coordinates of the landmarks to the “geomorph” R package to calculate distance (“interlmkdist” function; Adams et al., 2020). Due to the invasive and often lethal process of measuring the proboscis of individual bumblebees, we estimated prementum length (from the base of the mentum to the distal point on the basiglossal sclerite; Cariveau et al., 2016) using a caste- and species-specific allometric relationship with ITD from bees caught within a radius of 15 km from Mt Nuolja (Methods S2; Table S6-7; Fig. 2C). We used prementum length over total proboscis length (prementum plus glossa length), as the soft glossal tissue was easily damaged during the dissection process. To ensure the robustness of estimated prementum lengths, we excluded castes within species for which there were fewer than five observations for the scaling relationship (160 bees excluded, leaving 2,686). Repeatability of measuring bumblebee traits was >0.98 (Methods S3) and the conditional R^2^ of the allometric relationship was 0.911 (Fig. 2C).

**Figure 2:**
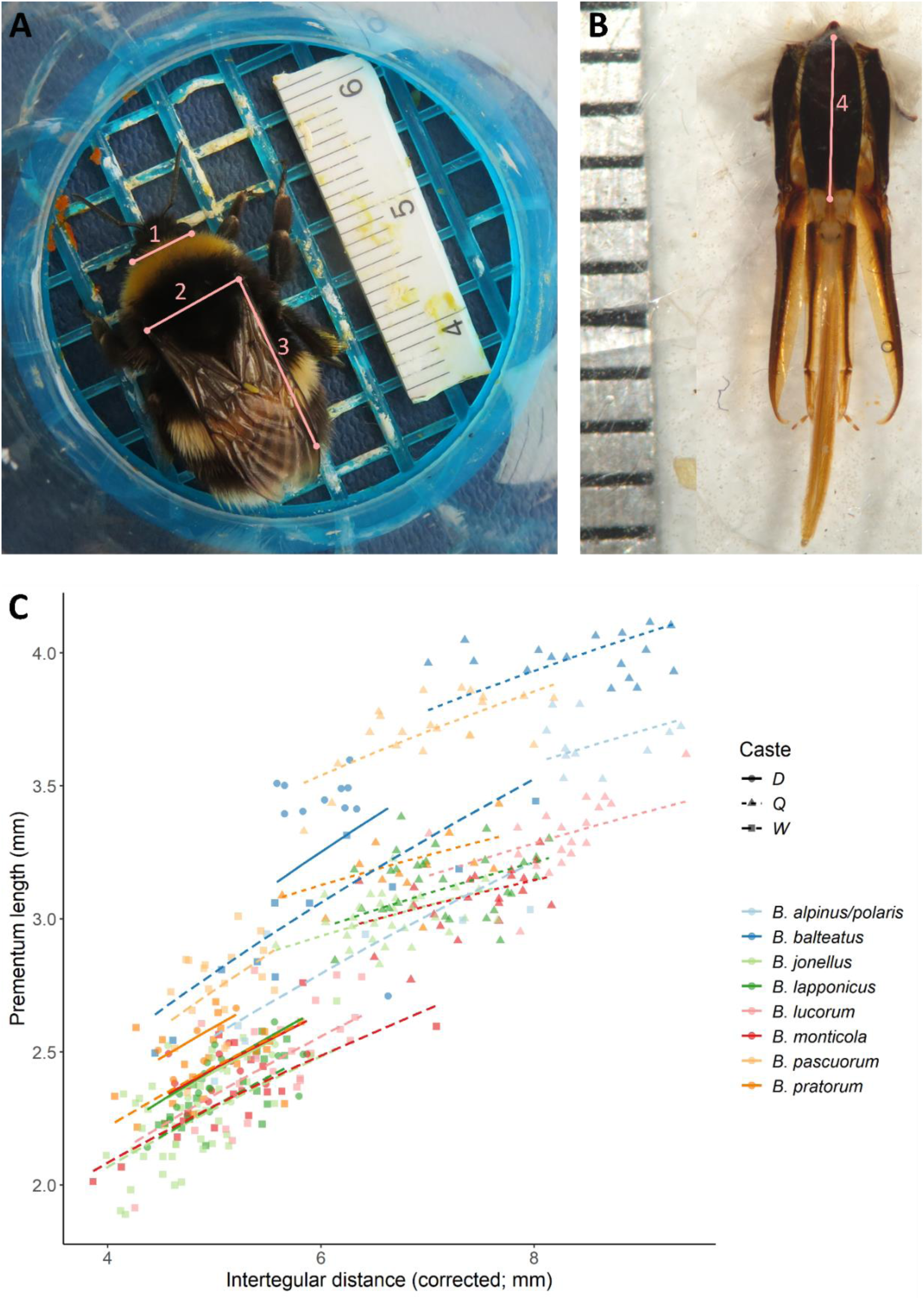
Bumblebee traits used to map bee trait space. A) We non-destructively measured head width (1), intertegular distance (2) and wing length (3) from images of bees (here, Bombus lucorum) caught during the bumblebee observation fieldwork (bees were released back into the wild after being imaged). B) We also measured bumblebee prementums (4) from lethally sampled bumblebees caught close to our field site to develop a scaling relationship between prementum length and ITD so that we could (C) estimate the prementum length of non-lethally caught bees during observation surveys.

We scaled and centred all continuous trait values (mean = 0; standard deviation = 1). To remove potential outliers, we excluded trait measurements that were >3 standard deviations from the mean of each trait for each caste nested within species (n = 19 individuals removed; Fig. S9). Finally, to reduce the number of trait axes and ensure each axis was orthogonal (required for hypervolume construction; see Section 2.4; Blonder et al., 2018), we performed dimensionality reduction of the continuous traits using PCA (using “prcomp” function from “stats” package; R Core Team, 2023) and took the first three axes (explaining 99.2% of the variation; see Fig. S10 for the contribution of each trait to each axis). We only included species with a minimum sample size of 20 (based on an analysis of simulated and real data; Methods S4; Fig. S11-13) and only included bees that had all four individual traits measured (n = 1,150). This left ten plant taxa: *Astragalus alpinus, Geranium sylvaticum, Melampyrum pratense, Pedicularis lapponica, Salix* spp., *Solidago virgaurea, Trollius europaeus, Vaccinium myrtillus, Vaccinium uliginosum* and *Vaccinium vitis-idaea*, visited by eight bumblebee taxa (22-304 visitations per plant taxon; Table S8; see Fig. S14 for the resulting interaction network; see Table S9 for our replication statement).

### 2.4 Building multidimensional trait space and statistical analysis

We constructed a hypervolume using the traits from every individual bee of all species to calculate a bumblebee community volume (Blonder et al., 2014, 2018). We built the hypervolume using the Gaussian kernel density estimation method (“hypervolume_gaussian” function from the “hypervolume” package; see Methods S4 for explanation of our choice and parameters; Blonder et al., 2018) and calculated its volume (using “get_volume” from the “hypervolume” package). Hypervolumes provide a useful framework for analysing multidimensional trait space, as they are flexible in the shape they take (Blonder et al., 2018). This ability to capture complex geometrical features is important, given the way plants interact with bee trait space can depend on a variety of factors, such as traits being important for overcoming exploitation barriers (Bartomeus et al., 2016), determining interaction efficiencies (Klumpers et al., 2019), or enabling environmental co-existence. Moreover, the relative importance of these factors may be species-specific.

#### 2.4.1 Do plant taxa exhibit trait space niches of the bumblebee community?

To examine, whether plant taxa interacted with specific regions of bumblebee community trait space, we first built a hypervolume for each plant taxon (again using the “hypervolume_gaussian” function), using only the bees that the specific plant taxon interacted with. We did this by down-sampling each plant taxon to 50% of its sample size 100 times (Lesser et al., 2020), constructing a hypervolume for each of these and then calculating its volume. Therefore, for each plant taxon, we had a distribution of values representing the volume of bumblebee trait space that it interacted with.

To gauge whether the volume of bee trait space that each plant taxon interacted with was smaller than would be expected if they interacted with bees at random, we used a randomisation procedure (Fig. S14). This randomisation procedure allowed a clear comparison between the volume of trait space used by each plant taxon compared to the volume available, and how this comparison could be affected by considering spatiotemporal constraints on bee trait space (Section 2.4.2). To do this, the plant taxon identity for each individual interaction was randomised (using “sample” from Base R; R Core Team, 2023) across the community 250 times. Following each randomisation, we built a new hypervolume in bumblebee trait space for each plant taxon and estimated its volume.

We then tested, for each plant taxon, whether the (distribution) volume of trait space they interacted with was significantly different from the corresponding null distribution. We did this by fitting a generalised least squares model (gls, with a Gaussian distribution; “nlme” package; Pinheiro et al., 2023) for each plant taxon, in which trait-space volume was predicted by the “volume type”, i.e. a categorical variable referring either to the null volumes or the actual volumes of bee trait space the plant taxon interacted with. As the distribution of null volume values and actual volume values differed in their variances, we included a “varIdent” variance structure in each model. To gauge how much smaller or larger the plant’s trait niche was relative to the null distribution (i.e. the effect size), we used the model-estimated contrast, and the associated standard error and (Bonferroni-corrected) *p*-value.

#### 2.4.2 How are our inferences of trait-space niches impacted when considering turnover?

We then evaluated the impact of trait turnover on the degree to which plant taxa interacted with a morphological subset of bee trait space. To do this, we repeated the randomisation procedure, but with the constraint that plant taxa could only interact with bees present during their 1) elevational ranges (spatial co-occurrence), 2) phenological range (temporal co-occurrence), or 3) elevational and phenological windows combined (spatiotemporal co-occurrence). These ranges were considered as continuous variables, i.e. elevation as meters above sea level and phenology as day of year. This meant that we compared again the volume (distribution) a given plant taxon interacted with against a distribution of “null” volumes, calculated by randomising the plant taxon identities present during the plant taxon’s elevational, phenological or spatiotemporal range. As plant taxa had different spatiotemporal ranges (Fig. 4), we performed this randomisation procedure for each taxon separately. We again tested for significance by fitting a gls for each plant taxon and gauged the effect size as the model-estimated contrast, following the same model structure and approach as in *2.4.1*.

Next, we tested whether changes in the degree to which plant taxa exhibited a trait niche (i.e. the effects sizes, or how much smaller or larger the plant taxa’s trait niches were relative to the randomised null values) was driven by phenological turnover or the elevational gradient. To do this, we statistically analysed whether these effect sizes differed among the turnover types considered. Therefore, we fitted a meta-analytic mixed-effects model (with Gaussian distribution, using the “rma.mv” function from the “metafor” package; Viechtbauer, 2010). In this model, the effect sizes were regressed against the “turnover type”, where turnover type refers to a dummy variable in the model that encodes whether the effect size was calculated when considering all trait space (ignoring turnover and plant-bee co-occurrence), or elevational, phenological or spatiotemporal turnover. As the effect sizes had been estimated from gls models, we propagated the uncertainty by weighting each effect size by its gls-estimated variance. Finally, as each plant taxon had four effect sizes in this model, we included plant taxon as a random intercept.

#### 2.4.3 How does turnover affect trait-space availability for each plant taxon?

To examine how each plant taxon’s spatiotemporal distribution affects the volume of bee trait space available to interact with, we compared 1) the actual trait space available (using bumblebees present across each plant taxon’s elevational range, phenological range, or spatiotemporal range) against 2) the total hypothetical trait space available (when the entire bee community is considered, regardless of whether they overlap in space and time with the plant taxon). To do this, we examined the total trait space volume (from *2.4*) against the volume recalculated using only bumblebees present across each plant taxon’s elevational ranges, phenological ranges, or spatiotemporal ranges. To reduce the impact of sample size on the estimations of hypervolume volume, we bootstrapped (100 times) each volume by subsampling the available bumblebee population to the smallest sample size (n = 75; using “sample” from Base R; R Core Team, 2023).

To statistically test how accounting for turnover affects the available trait space for each taxon, we fitted a generalised least squares model (with Gaussian distribution; “nlme” package; Pinheiro et al., 2023) in which the trait-space volume available (response variable) was regressed against the specific aspect of turnover (no turnover (or all bees), elevational, phenological or spatiotemporal). To test if bee trait-space availability differed significantly among the ten plant taxa and whether plant taxa were differentially affected by accounting for turnover, we included an interaction term between plant taxon and turnover type. As the plant taxa and turnover types showed heterogeneity of residual variance (due to the bootstrapping process), we included a variance covariate for the plant taxon and turnover type interaction (using the “varIdent” variance structure). We assessed the significance of each model term using likelihood ratio tests.

We conducted all analyses using R v. 4.3.2 (R Core Team, 2023).

## 3. RESULTS

### 3.1 Did plant taxa have trait niches of the bumblebee community?

#### 3.1.1 When all interactions are pooled regardless of occurrence in space and time (turnover not considered)

Plant taxa interacted with specific regions of bee trait space (Fig. 3), when compared against the trait space of the entire bumblebee community. This was evidenced by all plant taxa (except *Vaccinium myrtillus*) interacting with smaller volumes of bee trait space than the mean hypervolume volumes of the null distributions (i.e. having negative effect sizes; Fig. 5). Further, for eight of the ten plant taxa, this interaction with a morphological subset was significant (smaller hypervolume volumes than the null distribution: effect sizes ranged between −0.723 and −7.39, all *t*-values ≤ −4.42 & all *p* ≤ 0.01; Table S6; Fig. 6). In contrast, *Vaccinium myrtillus* interacted with a significantly larger volume of trait space than the randomised null distribution (effect size: 2.07; *t* = 14.7; *p* <0.001).

**Figure 3:**
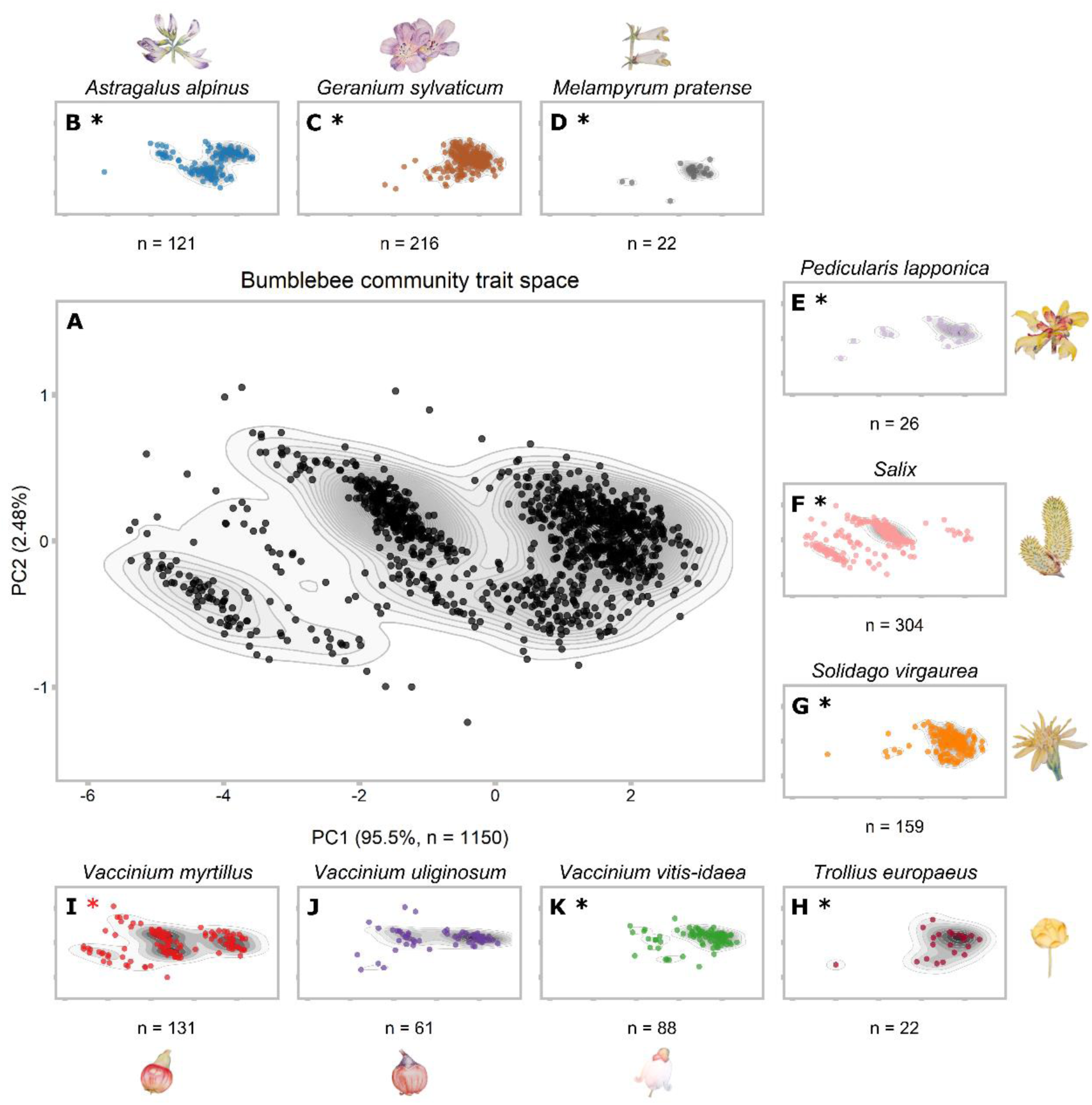
Bumblebee community trait space (A), and the bee trait space that each plant taxon interacted with (B-K). Each point represents an individual bumblebee sampled from eight taxa. Axes correspond to the first two dimensions of trait variation from a principal components analysis of four bumblebee morphological traits, with the first axis corresponding primarily to intertegular distance and the second to prementum length (Fig. S10). To see how plant taxa interacted with each PC axis independently, see Fig. S16. Black asterisks indicate species that we found significantly interacted with subset of bee trait space when turnover was not considered, whereas the red asterisk (3I) represents interacting with a significantly larger region of trait space than random. Drawings of plant taxa correspond to the subplot to which they are closest (not drawn to scale).

**Figure 4:**
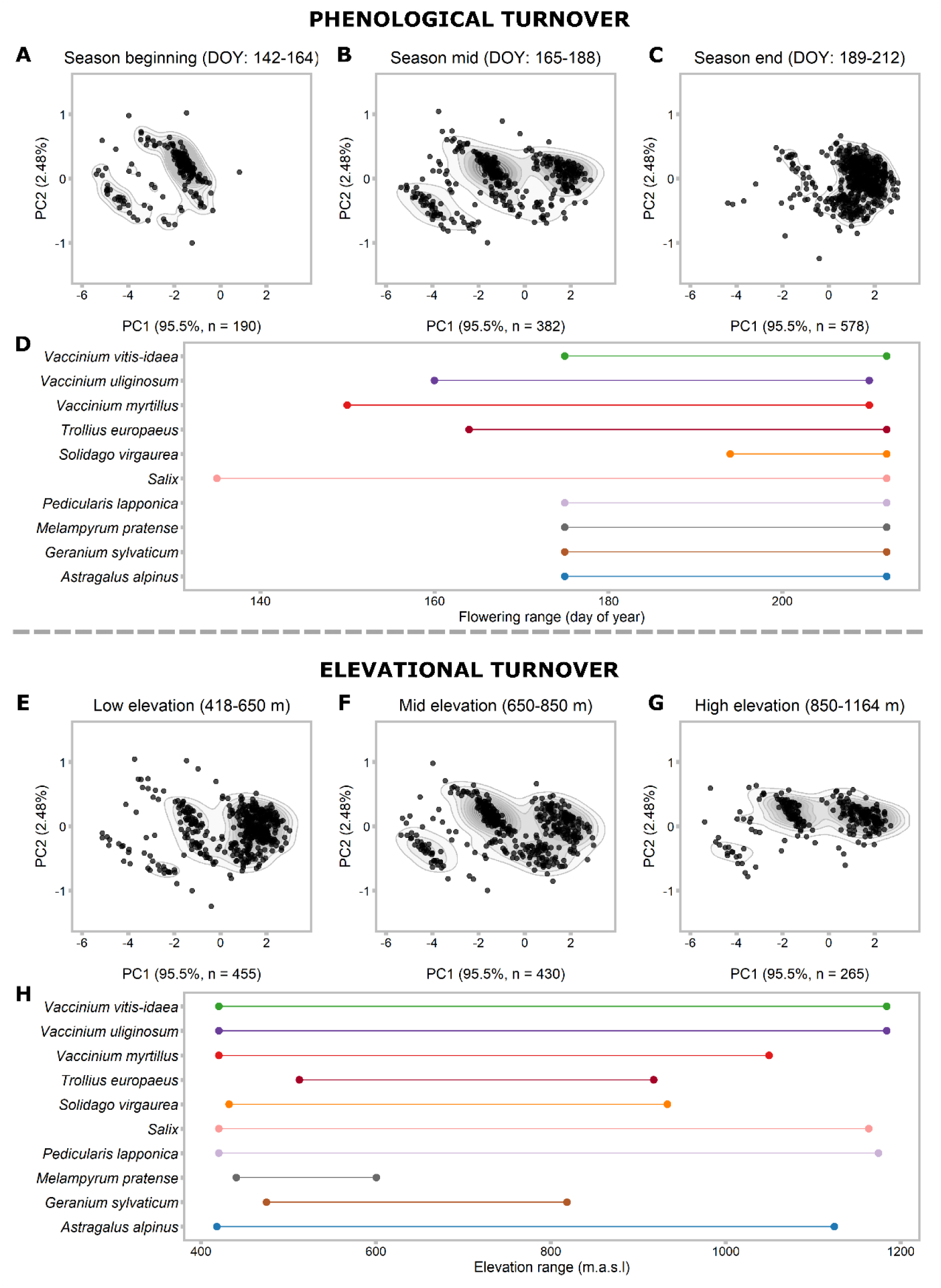
Turnover of bee trait space (A-C & E-G) and plant taxa (D & H) over phenological and elevational gradients. Phenological turnover of bee trait space is represented in two-dimensional space by splitting the season into three equally long sections: beginning (A; day of year [DOY]: 142-164); mid (B; DOY: 165-188); and end (C; DOY: 189-212). Spatial turnover of bee trait space is represented by splitting the elevational gradient into three sections, corresponding to the forest (E; vegetation zone A & B; low elevation: 418-650 m.a.s.l), willow (E; vegetation zone C; mid elevation: 650-850 m.a.s.l) and tundra zones (G; vegetation zones D & E; high elevation: 850-1164 m.a.s.l). Note, in the main analysis, we considered space and time as continuous variables. In A-C and E-G, each point represents an individual bee. Subplots D & H represent the phenological and elevational distributions of plant species, respectively. For visualisation, the phenological and elevational ranges show the lowest and highest values for the five years; however, in the main analysis, yearly ranges were used. For D, the y axis limit represents the end of our bee sampling period. For bee species distributions over the elevational gradient and season, see Figs. S5 & S6, respectively.

**Figure 5:**
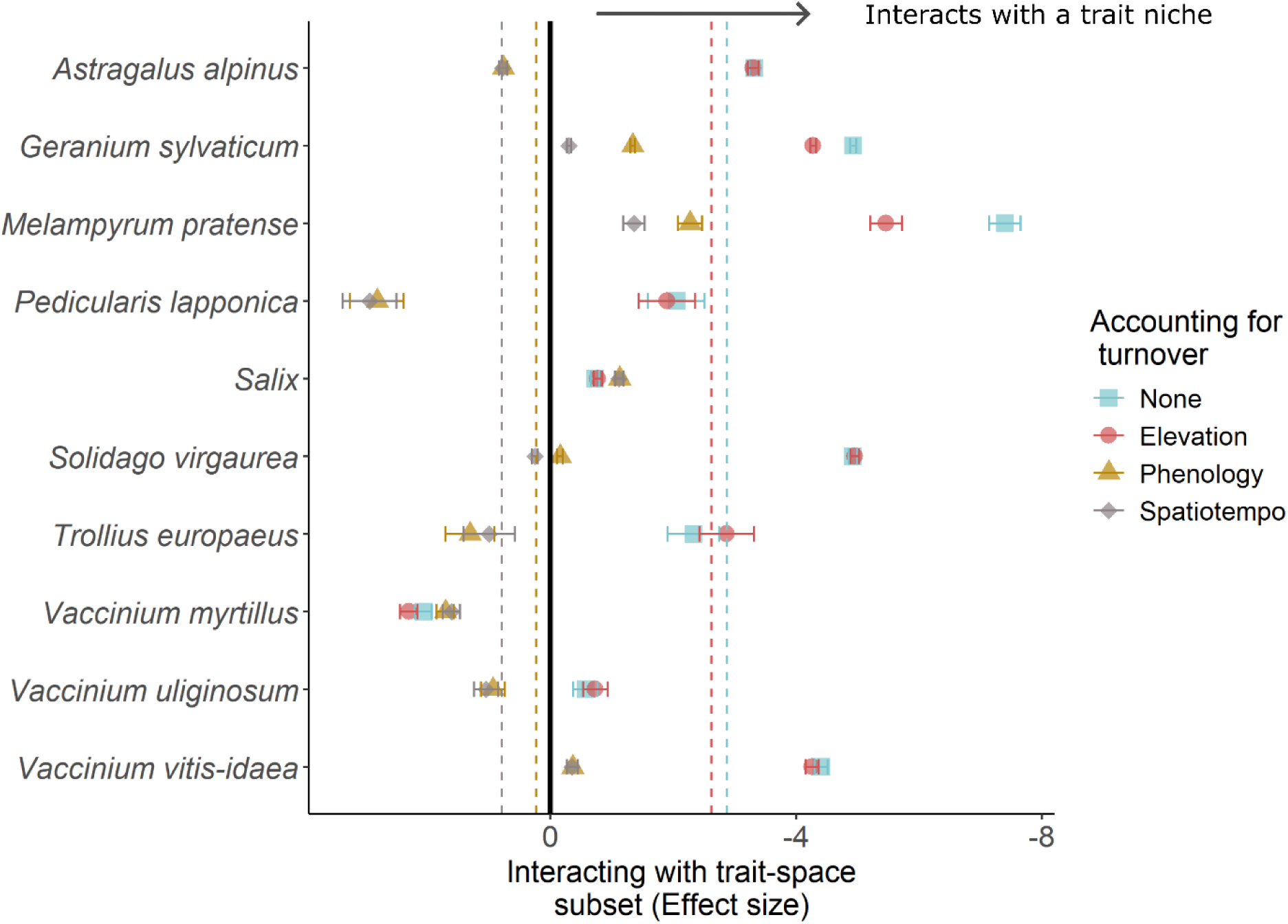
Accounting for different types of turnover (i.e. plant-bee phenological, elevational or spatiotemporal co-occurrence) impacts the degree to which plant taxa show a bee trait niche. Change in trait-niche effect size when plant taxa can interact with all bumblebee individuals in the randomisation (“None”), compared to when plant taxa can only interact with bees that co-occur in their elevational, phenological, or spatiotemporal ranges, represented as different coloured and shaped points. Error bars represent the standard error. An effect size close to zero (solid black line) indicates that a plant taxon interacted randomly with bees across the trait space available during their phenological, elevational or spatiotemporal range. More negative values (right of the solid line) indicate plant taxa that interacted with significantly smaller volumes of bee trait space than the randomisation, i.e. had a trait niche. The dashed horizontal lines represent the estimated mean effect sizes for each turnover type considered.

**Figure 6:**
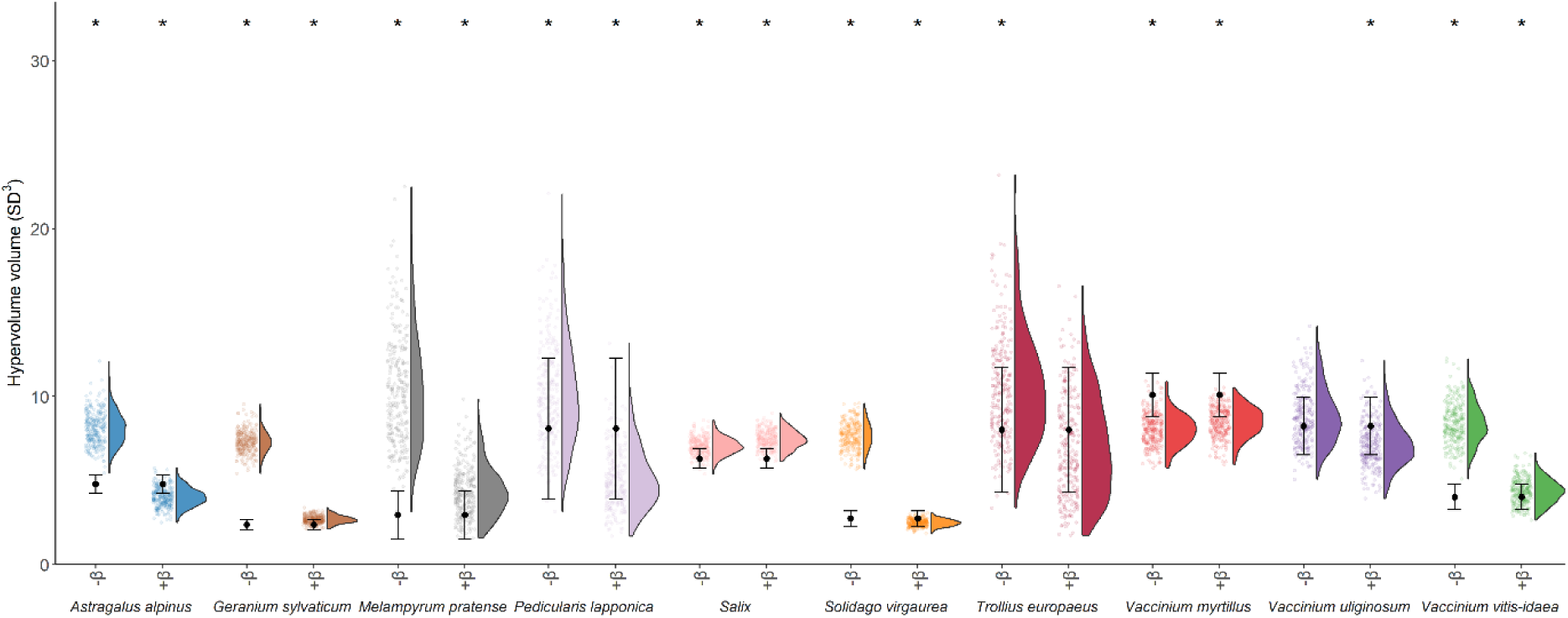
Change in degree to which plant taxa interacted with specific regions of bumblebee community morphological trait space (demonstrated as a trait niche), when turnover is not considered and when it is considered. Black points represent the observed volume of bumblebee trait space each plant taxon interacted with. Coloured distributions (and associated coloured points) represent the null expectations (with each point indicating a null value), based on random sampling of bees across either the entire bumblebee community trait space (“-β”), or only the trait space available during each plant taxon’s spatiotemporal distribution (“+β”). Asterisks indicate whether the volume of trait space that the plant taxon interacted with is significantly smaller than the null expectation, without or with accounting for turnover.

### 3.2 How much bumblebee trait space is available to plant taxa, when spatiotemporal turnover is considered?

Figure 4 shows that when we matched up plant spatiotemporal turnover with bee trait-space turnover, we can qualitatively observe that plant taxa co-occurred with different regions of bee trait space. Indeed, we found that the area of bee trait space available for plant taxa to interact with is significantly smaller (Fig. S17; likelihood ratio test (LRT): DF = 3, *F* = 19,486, *p* <0.001; Table S7) relative to when turnover is not considered (mean total bee trait space volume when turnover is not considered (8.62 SD^3^ (±1.43)). This was primarily driven by accounting for plant phenology, given the large trait turnover that occurs within bee species over time (Methods S1), as the mean trait space after accounting for temporal compared to spatiotemporal ranges differed little (5.69 SD^3^ (±2.25) versus 5.48 SD^3^ (±2.50), respectively). In contrast, there was almost no difference in available trait space after accounting for plant elevational ranges (8.26 SD^3^ (±1.94)). The degree to which trait space contracted was highly plant-taxon specific (LRT: DF = 27, *F* = 411, *p* <0.001), with *Salix* still having 100% of the total bumblebee trait space available even after accounting for its spatiotemporal distribution, compared to *Geranium sylvaticum*, for which bee trait space contracted to 3.17 SD^3^ (±0.640).

### 3.3 Did plant taxa still have a trait niche once plant taxon turnover and bee trait turnover were considered?

When only potentially feasible interactions were included (spatiotemporal turnover accounted for), the degree to which plant taxa still associated with specific regions of bee trait space depended on the type of co-occurrence considered (Fig. 5; Table S6). When only elevational co-occurrence was considered (red circles in Fig. 5), the degree to which plant taxa demonstrated a trait niche was slightly but significantly smaller than when turnover was not considered (blue squares in Fig. 5; from meta-analysis: estimated mean trait-niche effect size with no turnover = −2.87; mean effect size after accounting for elevation = −2.61; *z* = 6.13; *p* <0.001; Table S10 & 8). However, nine of the ten plant taxa still demonstrated a trait niche, after accounting for elevational turnover (gls: all effect sizes ≤ - 0.728; all *p* ≤0.0113).

However, when phenological (brown triangles in Fig. 5) or spatiotemporal (grey diamonds in Fig. 5) turnovers were considered, we found that most plant taxa no longer associated with specific regions of bee trait space (mean estimated effect size from meta-analysis: 0.231 & 0.790, respectively). Indeed, after accounting for spatiotemporal turnover, six plant taxa had effect sizes that were larger than the means of the null randomisations (with this being significant for five; effect sizes ≥0.254, *t* ≥5.33; *p* <0.001). The four taxa that continued to demonstrate a trait niche were: *Geranium sylvaticum*, *Melampyrum pratense*, *Salix* spp. and *Vaccinium vitis-idaea* (effect sizes ≤ −0.303, *t* ≤ - 4.02, *p* ≤ 0.003; Fig. 6; Table S10).

## 4. DISCUSSION

To assess how multidimensional bee trait space is associated with plant-bumblebee interactions, we characterised the trait niche of each plant taxon by calculating the area of trait space occupied by its interaction partners (individual bumblebees) (Dehling et al., 2016, 2022). We then investigated how appreciating the dynamic nature of this Arctic community – with turnover of species and traits – might impact the role of morphology in determining interactions. We therefore examined how trait-space availability changed when we accounted for bee trait turnover and each plant taxon’s spatiotemporal range, and whether accounting for this impacts our inference of multidimensional trait niches.

When we examined the network of plant-bumblebee interactions across the whole season and elevational range, most (eight of ten) plant taxa had trait niches (interacted with specific regions of bee trait space). However, when we considered the large spatiotemporal turnover in bumblebee trait space, the degree to which taxa exhibited trait niches declined leaving only fourplant taxa with bee trait niches. . By resolving localised dynamics of community turnover and linking them to individual-trait-based networks, our findings indicate that the ability of morphological traits to explain interactions might be spatiotemporally scale-dependent (González-Varo & Traveset, 2016; Klomberg et al., 2022) and taxon-specific (Junker et al., 2013). This scale dependency highlights the challenges in inferring mechanisms across spatiotemporal scales especially in highly dynamic systems like the Arctic when trying to forecast interaction rewiring.

### 4.1 Available multidimensional trait space is plant taxon-specific

When we compared the bumblebee trait space to what was actually available for a given plant taxon to interact with (co-occurrence), we found the position and volume of the available trait space differed among plant taxa. This was due to the differing phenological and elevation ranges of plant taxa (MacDougall et al., 2021) and the fact that bee trait space itself turned over (Cantwell-Jones et al., 2023). This meant that taxa that flowered for longer (such as *Salix*) retained access to greater regions of bee trait space than other plant taxa in the same network (although note that *Salix*’s phenology may be inflated due to being an aggregate of several species). Specifically, phenology contributed most to this taxon-specific trait space availability, rather than elevation range. Given bumblebees are social insects, this trend reflects the change in trait space that occurs when the actively foraging community switches from being dominated by queens (larger-bodied females) to workers (smaller-bodied females) (Cantwell-Jones et al., 2023; Methods S1). In contrast, the effect of accounting for elevation range was much smaller in magnitude. This result was surprising, as harsher abiotic conditions at higher elevations are often expected to constrain the morphologies, such as body size (Classen et al., 2017). However, this result may have been due to seven of the ten plant taxa being present along most of the elevational transect, meaning their elevational ranges did not limit the trait space available. Alternatively, although some of the bumblebee species have distinct elevational and habitat preferences (such as *B. pascuorum* being found primarily at low elevation; Fig. S5), individual bees can often use the entire elevational range on days with favourable abiotic conditions, especially later in the season (Lundberg & Ranta, 1980), potentially leading to transient increases in trait-space availability at higher elevations. Despite this, for the three taxa with more limited elevational ranges (*Geranium sylvaticum*, *Melampyrum pratense* and *Trollius europaeus*), accounting for elevation reduced trait space availability for *G. sylvaticum* and *M. pratense* (Fig. S17), suggesting that plant taxa with broad versus restricted elevation ranges do experience some difference in bee trait space availability, with which to interact.

This plant-taxon-specificity of bee trait space calls into question the scales at which we should be studying the mechanisms underpinning interactions, as the appropriate spatiotemporal scale may also be taxon-specific. For instance, it could be useful to consider pairwise interactions and/or trait space as being nested within the spatiotemporal ranges of the taxa within a community (as has been applied within food webs; Albert et al., 2023). This should help prevent “forbidden traits”, where two species do not interact due to compatible regions of trait space only being present outside of their spatiotemporal ranges (Olesen et al., 2011), when considering how trait space can be used to predict mutualistic interactions. Indeed, these forbidden traits could in part explain why species interactions can vary more across space than explained by species turnover (such as in host-parasite interactions; Poisot et al., 2017): matching trait combinations between interacting species may exist in one location in space and time but be absent in other locations due to trait turnover (Gómez et al., 2020; Klomberg et al., 2022).

Although we focused here on the bumblebee trait space available for plant taxa, it would also be interesting to extend this research to examine how plant trait space available for bumblebees changes spatiotemporally and how the relative abundances of different plant taxa (and their traits) shape trait niches of bees. Indeed, not only is there likely to be plant taxon turnover with elevation and over the season (MacDougall et al., 2021), but also changes to trait-space volume and position of the centroid. Much work has focused on how plant vegetative traits change with elevation (Kichenin et al., 2013; Rixen et al., 2022; Sundqvist et al., 2013), with the most conspicuous being plant height below and above the treeline (Mayor et al., 2017). Less work, however, has investigated how floral traits change with time or space (but see e.g. Gómez et al., 2020; Junker & Larue-Kontić, 2018). For example, floral colour has been observed to change with elevation (at the interspecific level: Bergamo et al., 2018; Gray et al., 2018; Shrestha et al., 2014), as has corolla length, nectar volume and nectar concentration (when measuring individuals across ten *Rhododendron* species; Basnett et al., 2019). It is therefore possible that alpine-specialist bumblebees might be exposed to a different plant trait space than subalpine species.

Additionally, although we focused on bumblebee pollinators to ensure feasible collection of high-resolution interaction- and trait-data, incorporating other pollinator species may increase trait-space turnover due to differences in phenology. For example, as bumblebees are the primary pollinators at the beginning of the season (Goulson, 2010; Kevan, 1973), it might be expected that trait space would expand as other pollinating taxa emerge, given the large trait differences between bumblebees and other Arctic pollinating taxa, like flies (Elberling & Olesen, 1996).

### 4.2 Plant taxon and bee trait turnover matter when determining multidimensional bee-trait niches at localised scales

When spatiotemporal turnover was not considered, nine of our ten plant taxa interacted with smaller volumes of bee trait space than the randomisation (and this was significant for eight). This morphological niche is in line with traditional research on plant-pollinator interactions (e.g. Stang et al., 2009), suggesting specific morphological traits could be important for overcoming exploitation barriers (i.e. proboscis longer than the corolla; Alexandersson & Johnson, 2002), determining interaction efficiency (Klumpers et al., 2019) or ecological fitting (Janzen, 1985). The lack of bumblebee trait niche for *Vaccinium myrtillus* is interesting, given its congener *V. vitis-idaea* had a trait niche (also a bell-shaped flower), highlighting the challenges that could come when using floral shape to predict pollinator trait niches (Peralta et al., 2024).

However, when we accounted for turnover in plant taxa and bee trait space (i.e. plant taxa could only interact with spatiotemporally available bees in the randomisation), the degree to which plant taxa had a bee trait niche declined and the number of taxa having a “niche” dropped from nine to four. This finding leads to two potential interpretations. Firstly, plant taxa might have evolved to co-occur spatiotemporally with the region of bee trait space that they are most compatible with, meaning their trait niches match the entire region of trait space that is spatiotemporally available (this may be especially true given we focused on bumblebee trait space out of the pollinator community). This is analogous to studies that find population-level floral morphology can covary with changes in the pollinator assemblage in space (Gómez et al., 2008; Izquierdo et al., 2023; Nagano et al., 2014) and time (Gómez et al., 2020).

Alternatively, plant taxa might simply be interacting with spatiotemporally available bees (e.g. CaraDonna et al., 2017), with the spatiotemporally available region of bee trait space having limited importance for determining interactions. CaraDonna et al. (2017), for example, found that the morphology of interacting Arctic plants and pollinators was poor at predicting weekly interaction rewiring in comparison to abundance, after phenology had been accounted for. Together with our findings, this could suggest that systems, like the Arctic, with short activity seasons and high turnover of species and traits could be composed of more generalist and opportunist species, meaning strategies to interact solely with specific morphotypes might be maladaptive. Indeed, previous work has found that trait-matching underpins plant-hummingbird interactions to a greater degree at lower latitudes and in regions with lower temperature seasonality (Sonne et al., 2020). Instead, in high turnover systems, spatiotemporal availability and abundance of plants or pollinators may be an overarching driver of interactions (Vázquez et al., 2007). This may especially be true for bumblebees, which can often be spatiotemporally flexible foragers (Sponsler et al., 2022). This explanation is additionally supported by our finding that *Geranium sylvaticum* and *Salix* spp were two of the taxa to continue demonstrating a trait niche when considering spatiotemporally available bees. Both taxa have relatively open floral morphologies, high abundance in our study site and high visual availability (i.e. raised from ground), potentially suggesting factors other than morphological barriers may be driving trait-space niches. Interestingly, although a longer phenology can enable more “generalist” interaction patterns (Bender et al., 2017; Glaum et al., 2021), we did not observe this for *Salix*.

These two possible explanations invoke different mechanisms underpinning plant-bumblebee interactions: trait matching versus abundance (Peralta et al., 2024). Disentangling the relative importance of each mechanism will be essential for forecasting how interactions might rewire under environmental change, as they suggest different levels of vulnerability for plant taxa in this system to persist or rewire (Peralta et al., 2020). For example, if plants have evolved to phenologically co-occur with their most compatible bumblebees, then phenological shifts under climate warming could lead to a mismatch between plant taxa and their most compatible pollinators (Duchenne et al., 2020; Hegland et al., 2009; Rafferty & Ives, 2012). However, if plants interact opportunistically with spatiotemporally available bees, they may be more able to rewire to new bumblebee partners over proximate timescales (Ponisio et al., 2017).

Although this study has given insight into how turnover can impact inferences of trait niches, there are still further questions that would be interesting to explore with more years of data. For example, to investigate how trait niches change over the elevational (or seasonal) gradient for plant species with large ranges. For example, Zhao et al. (2022) found dipteran pollinators exhibited trait matching at low elevation sites but not high, on a mountain in Southwest China. Assessing how trait space is differentially partitioned among plant species and the degree of overlap (for example, using multivariate analysis), and whether this changes spatiotemporally, would also be an interesting avenue for further research (Cantwell-Jones et al., 2024). Moreover, whilst the bee morphological traits chosen here (body size, prementum length, wing length, and head width) are generally considered important for plant-pollinator interactions (Eklöf et al., 2013), a myriad of other factors also likely underpin plant-bumblebee interactions, such as plant nectar and pollen quality (Klumpers et al., 2019; Somme et al., 2015; Vaudo et al., 2016), and interspecific competition (Brosi & Briggs, 2013). Ultimately, in our study system, the ability of morphological traits to predict interaction rewiring at small spatial scales and over proximate timeframes is not yet resolved. More broadly, we cannot accurately understand the mechanisms underpinning plant-pollinator interactions unless we incorporate turnover in species and traits.

### 4.3 Conclusions

By mapping turnover of individual bumblebee traits along an elevational gradient over the course of a season for five separate years, we found that each plant taxon experiences constraints on the traits it can interact with, due to spatiotemporal community turnover. When we considered these constraints, the degree to which plant taxa showed the expected trend of having a trait niche declined. To accurately understand the role of morphology in mediating plant-pollinator interactions, we therefore need to consider this spatiotemporal turnover, or we risk potentially overstating the importance of morphology when forecasting interactions under environmental change.

## Acknowledgements

ACJ is funded by the NERC Science and Solutions for a Changing Planet doctoral training programme, Imperial College London (NE/S007415/1). The project was also supported by two grants from INTERACT (funded by H2020 - agreement no. 730938, awarded to ACJ), funding from the Quekett Microscopical Club to RJG, the Geographic Field Grant from the Royal Geographic Society with IBG (to LH, SE and ACJ), the Heredity Field Grant from the Genetics Society (to ACJ), and the Fjällräven Field Grant from The Explorers Club to ACJ. We would like to thank T. Cox, C. Gibbons, R. Richardson, F. Brannlund and M. Aronsson for help with bumblebee sampling, and the Climate Impacts Research Centre (Umeå University) & Abisko Scientific Research Station for field equipment and their continued support. Finally, we would like to thank the Gill & Graystock laboratory groups for feedback on an early draft.

## Statement of authorship

ACJ, JE & RJG conceived the idea, with inspiration from OKB & JMT; KL initiated the phenology project; KL, OKB & RJG set up the permanent study site; ACJ, GA, LJ, MvU, SvB, MK, RL, JASJ, LB, SE, FC, LH, AMRAH, JS, OKB & RJG performed the fieldwork; ACJ measured morphological traits, with help from JE, JS & AMRAH; ACJ, JE & RJG developed the analytical approach, with feedback from JMT; ACJ performed the analyses with input from JE, JMT & RJG; ACJ & RJG wrote the manuscript, with feedback from all authors.

